# 40 Hz light stimulation restores early brain dynamics alterations and associative memory in Alzheimer’s disease model mice

**DOI:** 10.1101/2024.10.21.619392

**Authors:** Matthieu Aguilera, Chantal Mathis, Karin Herbeaux, Amine Isik, Davide Faranda, Demian Battaglia, Romain Goutagny

## Abstract

Visual gamma entrainment using sensory stimuli (vGENUS) is a promising non-invasive therapeutic approach for Alzheimer’s disease (AD), showing efficacy in improving memory function. However, its mechanisms of action remain poorly understood. Using young AppNL-F/MAPT double knock-in (dKI) mice, a model of early AD, we examined brain dynamics alterations before amyloid plaque onset. High-density EEG recordings and novel metrics from fields outside neuroscience were used to assess brain dynamics fluidity—a measure of the brain’s ability to transition between activity states. We revealed that dKI mice exhibit early, awake state-specific reductions in brain dynamics fluidity associated with cognitive deficits in complex memory tasks. Daily vGENUS sessions over two weeks restored brain dynamics fluidity and rescued memory deficits in dKI mice. Importantly, these effects built up during the stimulation protocol and persisted after stimulation ended, suggesting long-term modulation of brain function. Based on these results, we propose a “brain dynamics repair” mechanism for vGENUS that goes beyond current amyloid-centric hypotheses. This dual insight - that brain dynamics are both a target for repair and a potential diagnostic tool - provides new perspectives on early Alzheimer’s disease pathophysiology.

**Significance Statement:** Gamma ENtrainment Using Sensory stimuli (GENUS), involving 40 Hz rhythmic sensory stimulation, shows promise in improving memory function in Alzheimer’s disease (AD). We hypothesized that brain dynamics changes could be detected before plaque onset and modulated by vGENUS. Applying techniques from climate science to EEG recordings in young AD model mice, we found reduced brain dynamics fluidity associated with early cognitive deficits. Two weeks of vGENUS restored brain dynamics and improved memory, with effects persisting post-treatment. These findings challenge the amyloid-centric view of AD, introduce a potential early biomarker, and suggest vGENUS acts by “repairing” brain dynamics. Our approach offers new perspectives on early diagnosis and non-invasive interventions for AD and other neurological disorders with disrupted brain dynamics.

## Introduction

Alzheimer’s disease (AD) is a devastating neurodegenerative disorder and the leading cause of dementia worldwide, defined by the conjunction of progressive memory loss and cognitive decline with specific neuropathological changes. Despite its high and likely underestimated prevalence (1), effective treatments and early diagnostic tools remain elusive. Pathological hallmarks of AD include extracellular amyloid plaques formed by aggregated Amyloid-beta (Aβ) peptide, and intracellular neurofibrillary tangles of hyperphosphorylated Tau proteins. For decades, the amyloid hypothesis has dominated AD research, positing that Aβ accumulation and amyloid plaques induce network dysfunctions responsible for cognitive deficits (2) The diagnosis of AD requires the presence of both cognitive impairment and evidence of amyloid pathology which is detected either through cerebrospinal fluid (CSF) analysis via lumbar puncture to measure reduced soluble Aβ levels (3), or through neuroimaging techniques such as positron emission tomography (PET) to visualize insoluble Aβ deposits (4). However, mounting evidence challenges the timing and specificity of these diagnostic criteria: memory deficits often precede detectable amyloid plaque formation (5–8) and can occur independently of Aβ (9). Conversely, amyloid plaques are sometimes present in non-pathological or asymptomatic elderly individuals (10–12). While the amyloid pathway undeniably plays a role in AD (2), these findings underscore the need for alternative early biomarkers that capture other critical aspects of the disease progression.

One promising early indicator of AD, alongside memory deficits, is alterations in brain network activity, which can manifest before amyloid plaque onset (9). Advanced neuroimaging techniques such as functional Magnetic Resonance Imaging (fMRI) and Electroencephalography (EEG) enable the study of global brain dynamics, focusing on entire brain networks rather than region-specific changes. These tools have revealed that global brain dynamics follow scale-free patterns (13), slow down during aging (14), and are altered in AD (15–17). Thus, changes in global brain dynamics could serve as an early biomarker of AD, opening new windows for early diagnosis and intervention strategies.

The lack of effective treatments remains another major challenge in AD. While numerous drugs targeting Aβ reduction have been developed based on the amyloid hypothesis, many have failed to reverse AD symptoms and were discontinued due to side effects (18, 19). In this context, a novel non-invasive therapy based on 40 Hz gamma stimulation using sensory stimuli (GENUS (20)) has shown promising memory benefits in both AD mouse models (21, 22) and patients with mild probable AD (23) (for review, see (24)). Various mechanisms have been proposed to explain these effects, including reduced neurodegeneration and improved memory through lowered amyloid plaque load, primarily via microglial activation and glial responses (21, 22), or enhanced brain clearance (25).

However, recent studies have challenged these ideas, questioning the significance of gamma entrainment via sensory stimulations. Some report limited propagation beyond the visual cortex and lack of engagement in key regions such as hippocampal CA1 (26), while others debate whether true gamma oscillations are even being entrained (27). The impact of these stimulations on amyloid plaque loads also remains controversial (27, 28). Despite these ongoing debates, discussions about GENUS have largely remained focused on its effects related to the amyloid hypothesis.

We propose an alternative hypothesis: GENUS may restore memory by modulating global brain network dynamics, which are altered early in AD, rather than through 40 Hz-specific effects. This novel perspective is supported by recent evidence demonstrating GENUS can affect brain dynamics beyond AD pathology (29, 30), suggesting broader impacts on brain function.

To test this hypothesis, we conducted high-density EEG (hdEEG) recordings in a preclinical AD mouse model (AppNL-F/MAPT double knock-in mice (31); dKI, n=8, both sexes) and control littermates (n=8, both sexes) during memory task performance. Our previous work (32) demonstrated that at 4 months, dKI mice maintain normal performance in simple tasks (e.g., short-term Novel Object Recognition or Object Location) but exhibit specific and subtle deficits in more complex associative memory tasks such as Object-Place association. In this study, we evaluated memory performance and brain dynamics during the Object-Place association and long-term Novel Object Recognition tasks before and after 2 weeks of daily 1-hour visual GENUS (vGENUS) exposure.

## Results

### Young dKI mice display memory deficits in complex tasks before amyloid plaque onset

We first evaluate the extent of amyloid pathology in 4-month-old dKI mice. To this aim, we performed 6E10 immunohistochemistry on brain sections from 4 WT and 4 dKI mice (Fig. 1A, B; 2 males and 2 females per group). Across 262 brain slices from dKI mice, we detected only one plaque, in stark contrast to 12-month-old dKI positive controls (1 male, 1 female) where we observed more than 10 plaques per slice. These results confirm that 4-month-old dKI mice do not yet exhibit significant amyloid plaque deposition.

**Figure 1.**
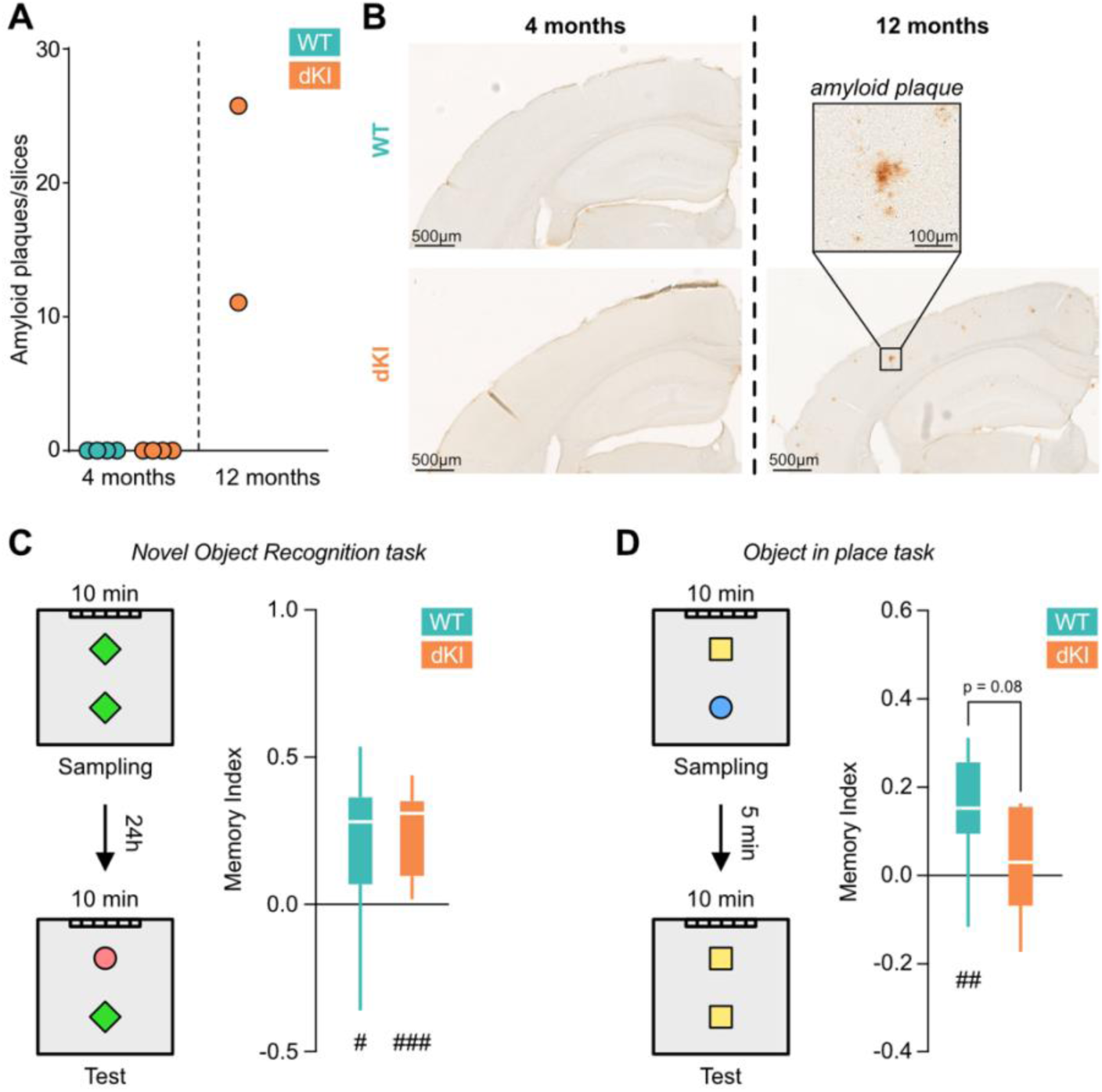
Young dKI mice display memory deficits in complex task before amyloid plaques onset. **(A)** Number of amyloid plaques per brain slices at 4 months (left) for WT (n = 4, blue) and dKI (n = 4, orange). We detect only 1 plaque in 262 brain slices from 4 dKI mice. As a positive control, plaques were counted for 12 months old (right) dKI (n = 2, orange). Plaques were counted over numerous frontal brain slices of several brain region plans. The total amount of plaques counted was divided by the total amount of slices for each animal. **(B)** Amyloid plaques immuno- staining of dorsal hippocampus at 4 months (left) for a WT (top) and a dKI (bottom) mouse and at 12 months (right) for a dKI mouse. Amyloid plaques were present in dKI mice at 12 months (top right for magnification) but not at 4 months. **(C)** Novel Object Recognition (NOR) task protocol (left), and memory index for WT (n = 8, blue) and dKI (n = 8, orange) mice (right). Both WT and dKI shows memory performances higher than chance levels (one-sided t-test against chance, #: p < 0.05, ##: p < 0.01, ###: p < 0.001). Box ranges from 25 to 75 percentile and whiskers for minimum to maximum values, median is represented by white line. **(D)** Object-in-Place (OiP) task protocol (left), and memory index for WT (n = 8, blue) and dKI (n = 8, orange) mice (right). dKI mices shows memory performances not higher than chance levels (one-sided t-test against chance, #: p < 0.05, ##: p < 0.01, ###: p < 0.001) and almost significantly lower than WT (two sample two sided t-test, p = 0.08). Box ranges from 25 to 75 percentile and whiskers for minimum to maximum values, median is represented by white line.

We next assessed the memory performance of dKI mice at this pre-plaque stage. The Novel Object Recognition (NOR) task with a 24-hour delay, a standard test for evaluating recognition memory (33), revealed no significant deficits in young dKI mice (Fig. 1C, SI Appendix, Fig. S1). This indicates that long-term recognition memory remains intact at this stage, unlike in mouse models with established amyloid pathology (34). However, when we employed a more complex memory task taxing short-term associative memory, the Object in Place (OiP) task, we detected subtle but significant early memory deficits in dKI mice (Fig. 1D, SI Appendix, Fig. S1).

These findings, consistent with our previous work (32), demonstrate that 4-month-old dKI mice exhibit measurable memory impairments in complex tasks prior to the development of significant amyloid plaque pathology. This provides a critical window to study early brain network alterations associated with AD and to test potential early interventions before substantial pathology develops.

### Global brain dynamics of dKI mice are altered before amyloid plaque onset

To investigate whether these early memory deficits were associated with alterations in global brain dynamics, we first focused our analysis on the OiP task, where we observed cognitive impairments. We recorded high-density EEG during the performance of the task and characterized global brain dynamics using an unbiased approach. This method treated multivariate EEG time-series as trajectories in a high-dimensional space of brain activity topographies (30 dimensions, corresponding to the number of hdEEG channels). Using t-SNE (35), a nonlinear, distance-preserving embedding method, we visualized these configurations as points in a two-dimensional space. For each point, we computed local dynamics fluidity (36), a metric related to the time the system takes to leave a neighborhood of the visited point in the space of dynamic configurations (Methods). This quantity, grounded in concepts from the statistical theory of extreme events (37) and previously used in the analysis of climatic time-series (36), offers important advantages on more classic metrics (38) of dynamic stability as it can be properly estimated from a considerably lower amount of data. While the overall shape of the sampled manifold was similar across genotypes, the average distribution of dynamics fluidity across the manifold was significantly lower in dKI mice (Fig. 2C Top; KS = 0.2283 ± 0.0277, p < 0.001). This reduction was associated with dKI mice exploring dynamic configurations not observed in WT mice, potentially corresponding to states with abnormally low fluidity (Fig. 2B, Top, where configurations unique to dKI mice appear as black points in the WT scatter plot). These findings suggest that by 4 months of age, dKI mice experience events where brain dynamics transiently exhibit reduced fluidity.

**Figure 2.**
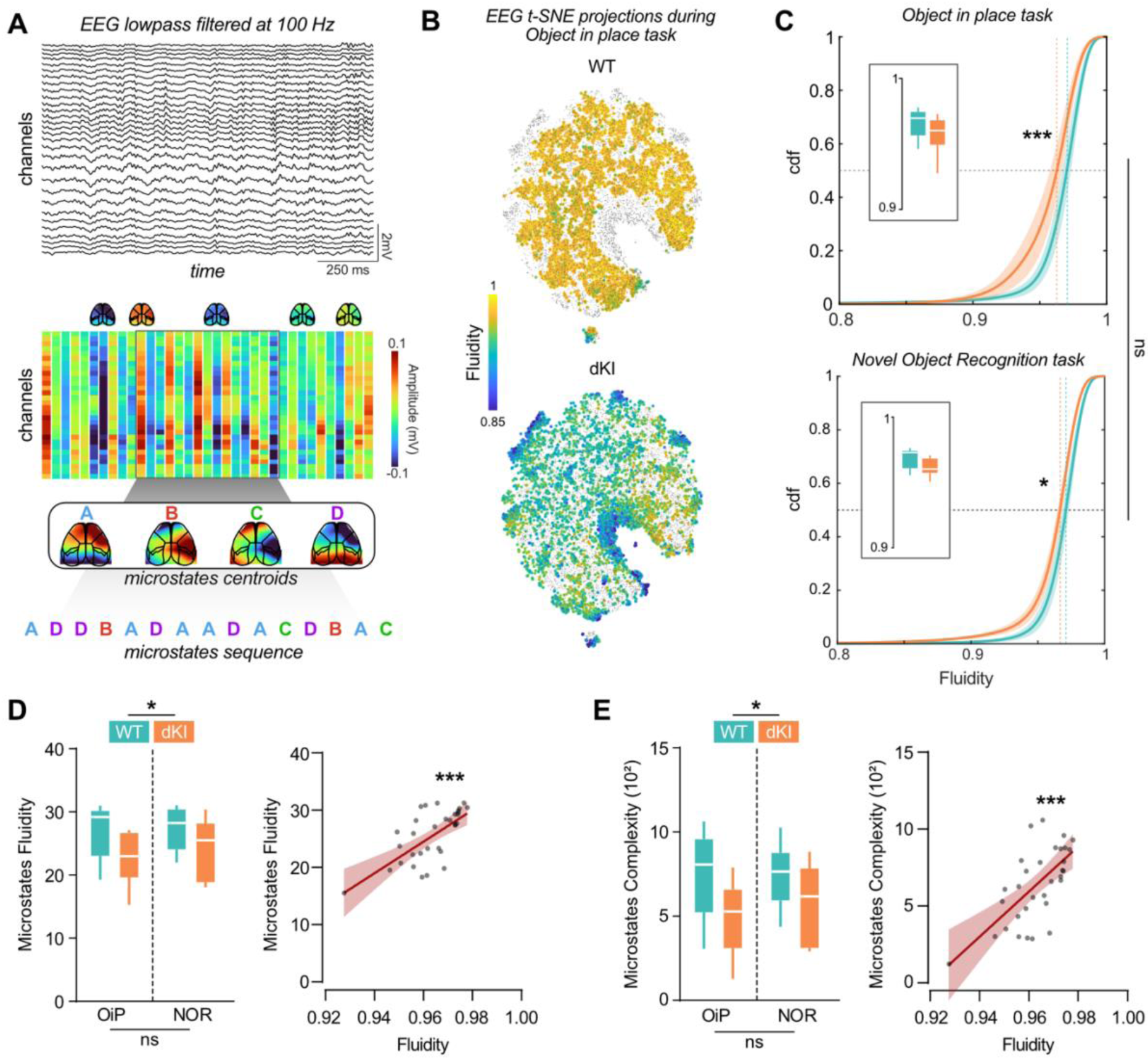
Global brain dynamics of dKI mice is altered before amyloid plaques onset. **(A)** Top, example of 30-channel hdEEG recorded during behavior. Middle, hdEEG is then coarsegrained with 40 ms window generating 30-channel amplitude vectors over time. Bottom, k- means clustering of these vectors identifies microstate centroids and sequences. **(B)** t-SNE projection of these previous vectors from a representative WT and dKI mouse during OiP task, color-coded by EEG dynamics fluidity. Grey dots represent the full distribution of both genotypes. **(C)** A shift between the cumulative distribution functions (cdf) of the EEG dynamics fluidity distributions for WT (blue) and dKI (orange) mice during OiP (top) and NOR (bottom) task (n = 8 per group) indicated reduced dynamics fluidity in dKI mice in OiP (KS = 0.2283 ± 0.0277, p < 0.001, here and for all other KS statistics in the following measured from a bootstrap comparison between association and null hypothesis, see Methods), and NOR (KS = 0.1626 ± 0.0279, p = 0.026). Data are presented as mean ± s.e.m. Dotted lines show distribution medians. Box displays individuals mean dynamics fluidity distributions. **(D)** Microstates fluidity during OiP and NOR task for WT and dKI mice (left, n = 8 per group). Two-way ANOVA indicate only a Genotype effect (F(1, 28) = 6.656, p = 0.015) indicating a reduced microstates fluidity in dKI mice for the 2 tasks. Box ranges from 25 to 75 percentile and whiskers for minimum to maximum values, median is represented by white line. Microstates fluidity correlates with mean dynamics fluidity (right, R² = 0.4539, p < 0.001). **(E)** Microstates sequence complexity during OiP and NOR task for WT and dKI mice (left, n = 8 per group). Two-way ANOVA indicate only a Genotype effect (F(1, 28) = 7.522, p = 0.011) indicating a reduced microstates sequence complexity in dKI mice for the 2 tasks. Box ranges from 25 to 75 percentile and whiskers for minimum to maximum values, median is represented by white line. Microstates fluidity correlates with mean dynamics fluidity (right, R² = 0.4371, p < 0.001).

To determine whether these alterations in brain dynamics were specific to tasks showing cognitive deficits, we also analyzed dynamics fluidity during the NOR task performance. Notably, dynamics fluidity was also significantly lower in dKI (Fig. 2C, Bottom; KS = 0.1626 ± 0.0279, p = 0.026), despite the absence of observable memory deficits in this task. We found no significant differences in dynamics fluidity between the two tasks for either WT (KS = 0.0558 ± 0.026, p = 1) or dKI mice (KS = 0.1296 ± 0.0263, p = 0.1468). This suggests that reduced dynamics fluidity is a general feature of dKI mice, independent of task-specific demands, and may serve as a sensitive early indicator of AD-related brain changes.

To corroborate these findings, we conducted EEG microstate analyses (39). Microstates represent a small number of stereotypical hdEEG topographies, extracted via unsupervised clustering. Continuous recordings were then converted into sequences of symbolic labels indicating the microstate closest to the current topography (Fig. 2A, Bottom). From these sequences, we calculated a fluidity-equivalent measure by quantifying the inverse probability of remaining in the same microstate from time t to t+1 (Methods). This measure was computed for several microstates (from 3 to 8) and then averaged to generate a single value. Microstates fluidity was significantly reduced in dKI mice in both OiP and NOR tasks compared to WT (Fig. 2D, Two-way ANOVA, F(1,28) = 6.656, p = 0.015) and showed a strong correlation with the previously computed dynamics fluidity (Fig. 2D, R² = 0.4539, p < 0.001).

Additionally, we assessed the complexity of microstate sequences using a minimum description length approach, which has previously been shown to be affected in AD (17). Consistent with the microstates fluidity results, microstate sequence complexity was significantly reduced in dKI mice during both OiP and NOR tasks compared to WT mice (Fig. 2E, Two-way ANOVA, F(1,28) = 7.522, p = 0.011) and also correlated strongly with dynamics fluidity (Fig. 2E R² = 0.4371, p < 0.001). These results were robust, remaining independent of the average number of microstates extracted (SI Appendix, Fig. S2), microstates self-repetitions (SI Appendix Fig. S2), and were confirmed using an alternative EEG microstates extraction method (SI Appendix Fig. S3).

### Global brain dynamics of dKI mice are unaltered during sleep

Given that global brain dynamics were altered in both OiP and NOR tasks, yet dKI mice showed memory deficits only in the OiP task, we sought to understand this discrepancy. We hypothesized that the difference might be attributed to either the complexity of the task (NOR being simpler than OiP) or the difference in delay between tasks. The 24h delay in the NOR task allows for sleep during the intertrial interval, potentially enabling offline memory consolidation. To address whether brain dynamics are altered during sleep, we analyzed dynamics fluidity during rapid eyes movements (REM) and slow-wave sleep (SWS) in the hours following learning in the 24h NOR task (Fig. 3A), as these states are critical for memory consolidation (40, 41).

**Figure 3.**
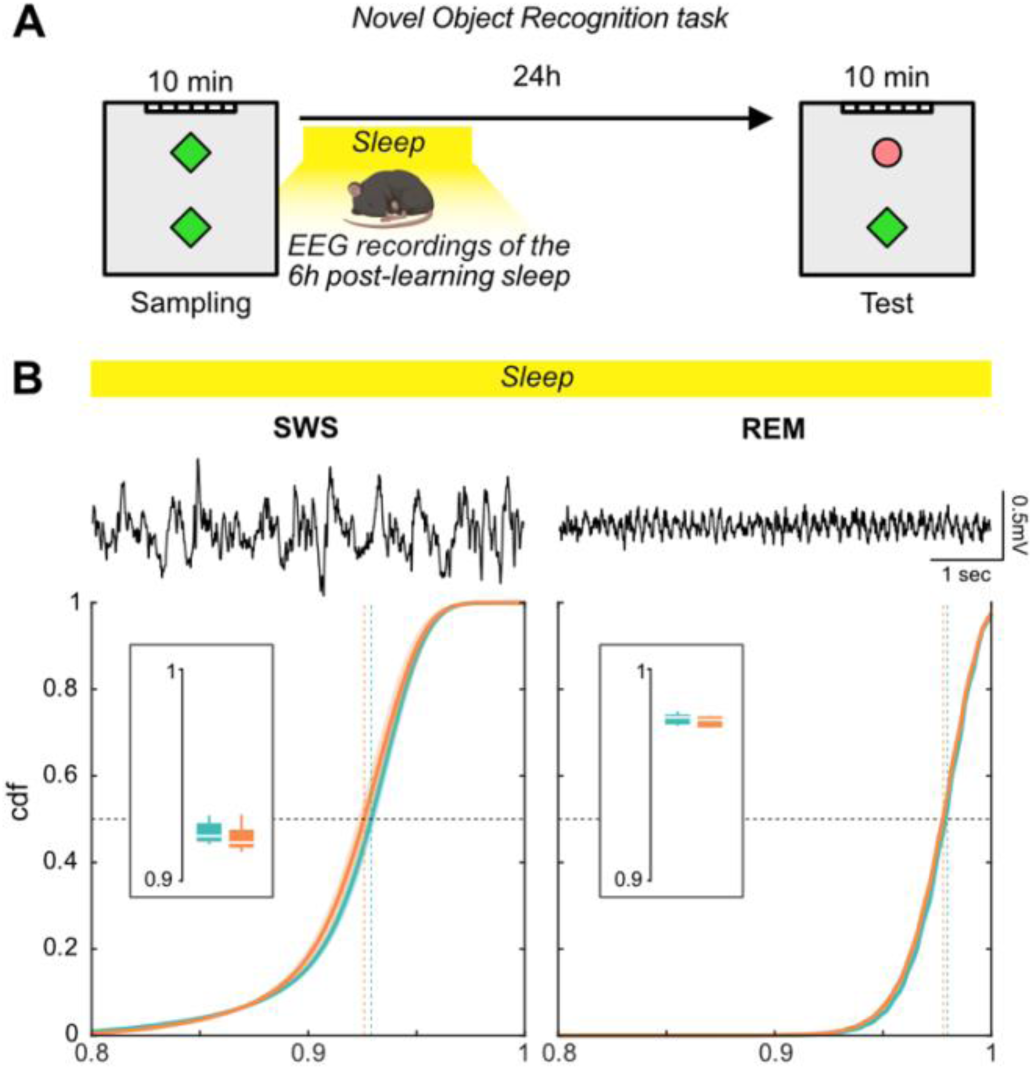
Global brain dynamics of dKI mice is unaltered during sleep. **(A)** EEG was recorded during the first 6 hours post-learning of the NOR task. **(B)** Top, example EEG signals during slow- wave sleep (SWS, left) and REM sleep (right). Bottom, no significant differences in EEG dynamics fluidity between WT and dKI mice during SWS or REM. Data are presented as mean ± s.e.m. Dotted lines show distribution medians. Box displays individuals mean dynamics fluidity distributions.

Intriguingly, we found that both SWS (KS = 0.0876 ± 0.0273, p = 0.3122) and REM (KS = 0.0653 ± 0.0232, p = 0.617) exhibited similar dynamics fluidity between dKI and WT mice (Fig. 3B). This preserved dynamics fluidity during sleep, in contrast to the altered dynamics during wakefulness, may explain the differential impact on behavior observed between the two memory tasks. Specifically, while the OiP task relies on online associative processes that may be disrupted by transient reductions in dynamics fluidity, the 24h NOR task may benefit from offline memory consolidation during sleep.

The intact dynamics during sleep might therefore help compensate for errors introduced by altered dynamics during wakefulness.

### vGENUS differentially entrains cortical regions at 40Hz and increases brain dynamics in dKI mice

Given the reduced dynamics fluidity observed in dKI mice, we next investigated whether vGENUS could potentially modulate these altered brain dynamics. We conducted 2 weeks of daily 1-hour vGENUS sessions and analyzed hdEEG for 10 minutes during stimulation on both the first and last (15th) day of the protocol to assess the immediate and long-term effects of the intervention. First, we examined the effects of vGENUS on cortical activity. EEG channels were grouped per cortical region based on Allen institute mouse brain parcellation to facilitate interpretation (Fig. 4A). A 40Hz peak in the power spectrum was observed across regions for both genotypes, with a higher proportion of the power spectrum in occipital (i.e. visual) regions (Fig. 4B, C). However, the relative 40Hz power during stimulation differed between genotypes and between Day 1 and Day 15 of the stimulation protocol (Fig. 4D, Three way repeated ANOVA, F_(1,84)_ = 4.619, p = 0.034). Specifically, relative 40Hz power during stimulation was lower in dKI mice at Day 1 (t(84) = 3.265, p = 0.009) but not significantly different from WT levels at Day 15 (t(84) = 1.975, p = 0.309). These findings suggest that the cortical response to 40Hz visual stimulation is initially reduced in young dKI mice but changes over the course of the 15-day vGENUS protocol.

**Figure 4.**
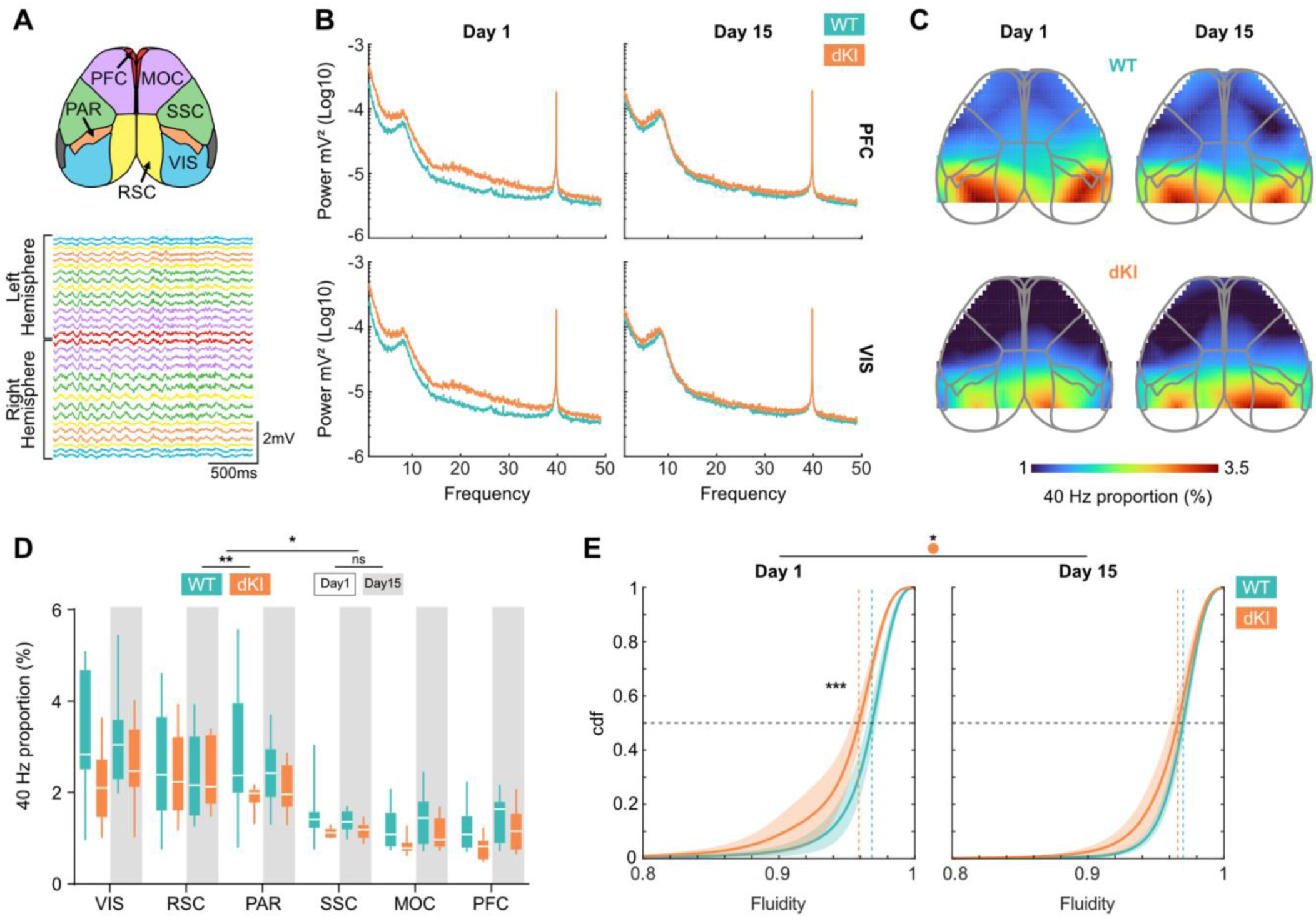
vGENUS differentially entrain cortical regions at 40Hz and increase brain dynamic in dKI mice. **(A)** Cortical parcelation map *(Top)* of the 30 channels hdEEG *(Bottom)* based on Allen Atlas, allowing the parcelation of the data into these following cortical regions: Visual Cortex (VIS, blue); Retrosplenial Cortex (RSC, yellow); Parietal Cortex (PAR, orange); Somato Sensory Cortex (SSC, green); Motor cortex (MOC, purple); Prefrontal Cortex (PFC, red). The power of these channels are averaged over regions for next analyses. **(B)** Power spectrum of PFC *(Top)* and VIS *(Bottom)* during first *(left)* and last *(right)* day of 40Hz vGENUS stimulation for one WT (blue) and one dKI (orange) mouse. A peak at 40Hz can be observed in both mice, both regions and both first and final day meaning a cortical entrainment by the vGENUS stimulation. **(C)** Cortical map of 40Hz proportion of the power spectrum (40Hz Power / all Power Spectrum) during first (left) and last (right) day of stimulation for one WT (Top) and one dKI (Bottom) mouse. As expected visual areas are the more entrained by 40Hz, this entrainment appears lower in the dKI mouse. **(D)** To quantify observations of previous maps, we measured the average 40Hz proportion of the power spectrum across different cortical regions during the first (no background) and last (grey background) day of stimulation for WT (n=8, blue) and dKI (n=8, orange) mice. Two-way ANOVA on repeated measures revealed a significant interaction between the day of stimulation and genotype (F(1,84) = 4.619, p=0.034). Post hoc tests showed that the 40Hz proportion was lower in dKI mice on Day 1 (t(84) = 3.265, p = 0.009), and no difference between genotypes on Day 15 (t(84) = 1.975, p =0.309). Box ranges from 25 to 75 percentile and whiskers for minimum to maximum values, median is represented by white line. **(E)** EEG Fluidity distribution during first (Left) and last (Right) day of vGENUS stimulation for WT (blue, n=8) and dKI (orange, n= 8) mice showed a lower EEG fluidity at Day 1 for dKI mice (KS = 0.2548 ± 0.0284; p < 0.001) which normalized to WT levels at Day 15 (KS = 0.1459 ± 0.0266; p = 0.066) after a specific increased fluidity in dKI mice (KS = 0.1587 ± 0.0275; p = 0.03). Data are presented as mean ± s.e.m. Dotted lines show distribution medians.

Given the ongoing debate on the significance of 40Hz stimulation, we next assessed whether vGENUS impacted global brain dynamics beyond its effects on 40Hz power. To this end, we computed dynamics fluidity during the stimulation periods on both Day 1 and Day 15. On Day 1, dKI mice exhibited significantly lower dynamics fluidity compared to WT mice (Fig. 4E, KS = 0.2548 ± 0.0284, p < 0.001). However, after 15 days of stimulation, dynamics fluidity in dKI mice increased (KS = 0.1587 ± 0.0275, p = 0.03), reaching levels comparable to WT mice (KS = 0.1459 ± 0.0266, p = 0.066).

These results suggest that while vGENUS has an immediate differential effect on 40Hz power in dKI mice, its impact on global brain dynamics becomes evident only after prolonged stimulation. This highlights the importance of chronic stimulation in promoting vGENUS effects on overall brain function and suggests that the mechanisms underlying these effects may involve processes beyond simple entrainment of cortical oscillations.

### Increased brain dynamics in dKI mice persist after end of stimulation and restore memory performance

Since memory deficits in young dKI mice were likely linked to altered brain dynamics, we examined whether vGENUS-mediated restoration of brain dynamics would also improve memory performance. At the end of the 2 weeks of vGENUS, NOR and OiP tasks were performed in the same order as before vGENUS (Fig. 5A). vGENUS did not affect performance in the NOR task, as dKI mice showed no memory impairments both before and after vGENUS (Fig. 5B, Left, SI Appendix Fig. S4). However, there was no longer a significant difference in dynamics fluidity between genotypes (Fig. 5B, Right, KS = 0.0798 ± 0.0235, p = 0.6405) due to a significant increase in dynamics fluidity in dKI mice after vGENUS (KS = 0.2137 ± 0.0293, p = 0.0042).

**Figure 5.**
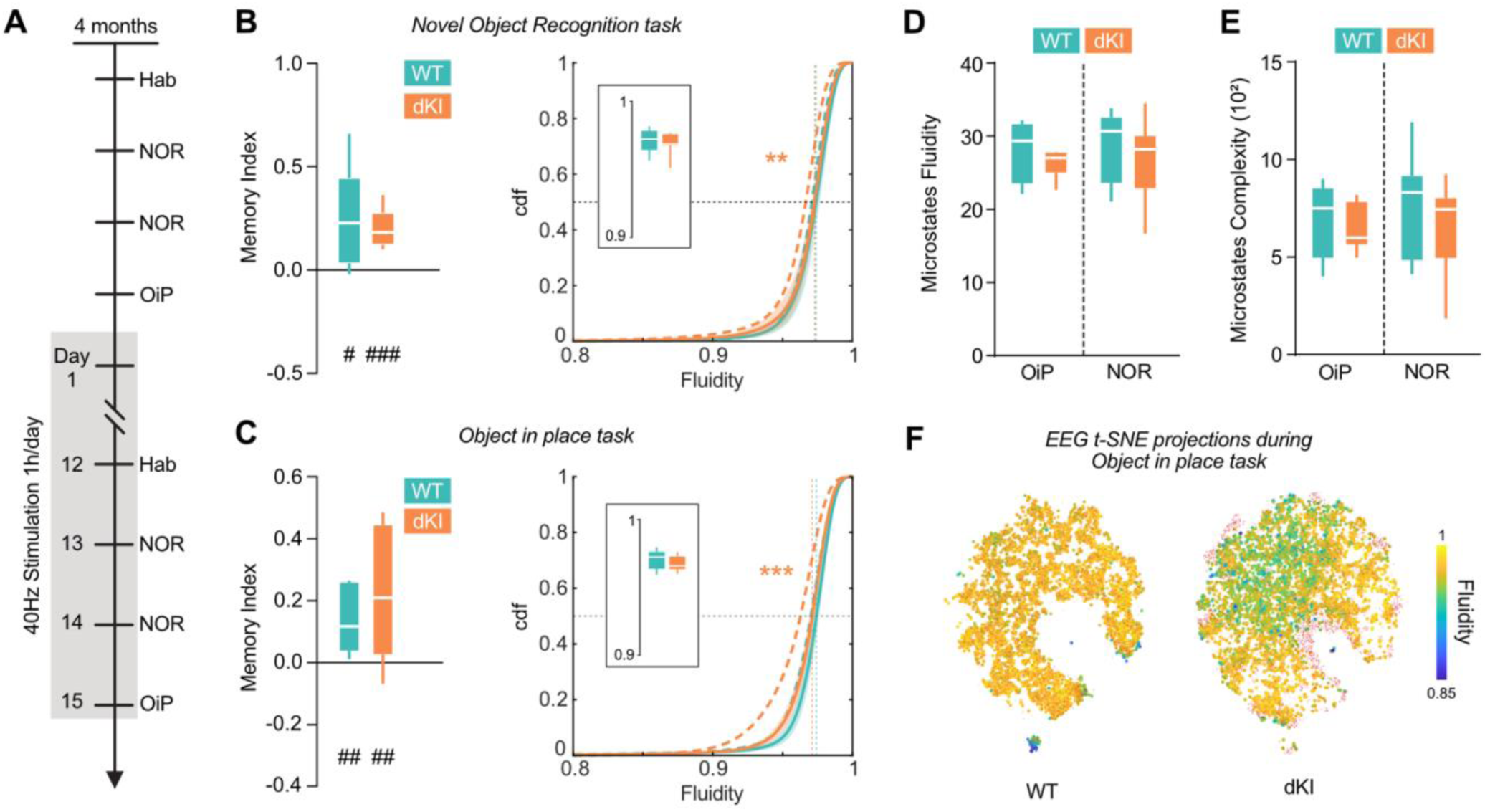
Increased brain dynamics in dKI mice remain after end of stimulation and restore memory performances in dKI mice. **(A)** 15 days vGENUS protocol schedule. **(B)** Post-GENUS NOR memory index for WT (n = 8, blue) and dKI (n = 8, orange) mice (left). Both WT and dKI shows memory performances higher than chance levels (one-sided t-test against chance, #: p < 0.05, ##: p < 0.01, ###: p < 0.001). Box ranges from 25 to 75 percentile and whiskers for minimum to maximum values, median is represented by white line. Brain dynamics fluidity show no more genotype differences (right, KS = 0.0798 ± 0.0235, p = 0.6405) due to a significant increase in dynamics fluidity in dKI mice after vGENUS (KS = 0.2137 ± 0.0293, p = 0.0042). Data are presented as mean ± s.e.m. Dotted lines show distribution medians. Box displays individuals mean dynamics fluidity distributions. **(C)** Post-GENUS OiP memory index for WT (n = 8, blue) and dKI (n = 8, orange) mice (left). Both WT and dKI shows memory performances higher than chance levels (one- sided t-test against chance, #: p < 0.05, ##: p < 0.01, ###: p < 0.001). Box ranges from 25 to 75 percentile and whiskers for minimum to maximum values, median is represented by white line. Brain dynamics fluidity show no more genotype differences (right, KS = 0.1298 ± 0.0270, p = 0.0999) due to a specific increase in dKI mice (KS = 0.237 ± 0.0283, p < 0.001) after vGENUS. Data are presented as mean ± s.e.m. Dotted lines show distribution medians. Box displays individuals mean dynamics fluidity distributions. **(D)** Microstates fluidity during OiP and NOR task for WT and dKI mice post vGENUS (n = 8 per group). Box ranges from 25 to 75 percentile and whiskers for minimum to maximum values, median is represented by white line. **(E)** Microstates sequence complexity during OiP and NOR task for WT and dKI mice post vGENUS (n = 8 per group). Box ranges from 25 to 75 percentile and whiskers for minimum to maximum values, median is represented by white line. **(F)** t-SNE projection of hdEEG multidimensional vectors from a representative WT and dKI mouse during OiP task after vGENUS, color-coded by EEG dynamics fluidity. Red dots represent the full genotype distribution pre and post vGENUS combined, highlighting genotype-specific patterns.

Thus, the vGENUS-induced increase in brain dynamics appears to persist even after stimulation sessions, suggesting a stable shift toward more physiological brain dynamics that could benefit cognitive performance. Indeed, after two weeks of vGENUS, dKI mice no longer showed deficits in the OiP task (Fig. 5C, Left, SI Appendix Fig. S4). Similarly to the NOR task, brain dynamics fluidity during the OiP task showed no significant difference between genotypes after vGENUS (Fig. 5C, Right, KS = 0.1298 ± 0.0270, p = 0.0999) due to a specific increase in dKI mice (KS = 0.237 ± 0.0283, p < 0.001).

Discretized microstates analysis supported these findings, showing no significant difference between genotypes in either microstates fluidity (Fig. 5E, SI Appendix Fig. S5) or microstates sequence complexity (Fig. 5F, SI Appendix Fig. S5) after two weeks of vGENUS.

Our results demonstrate that vGENUS restores memory performance in the OiP task by significantly increasing dynamics fluidity, even after the end of the stimulation, specifically in dKI mice. Indeed, vGENUS appears to specifically target the previously identified states of low dynamics fluidity in dKI mice. This effect can be visualized in Fig. 5D, where configurations present in dKI mice before vGENUS but absent after treatment are shown as red points. By eliminating these low fluidity states, vGENUS effectively “thaws” brain dynamics, returning them to a more fluid state similar to that observed in WT mice. Importantly, these effects of vGENUS on brain dynamics fluidity appear to be specific to pathological states. Sleep dynamics, which were unaffected in dKI mice before treatment, showed no differences in fluidity after vGENUS (SI Appendix Fig. S6), suggesting that vGENUS selectively targets abnormal awake brain states without disrupting normal sleep dynamics.

## Discussion

Using high-density EEG recordings during specific memory task performances in a preclinical AD mouse model and assessing global brain dynamics through novel metrics originally employed in fields outside of neuroscience (36), we have demonstrated that dKI mice exhibit early, awake state-specific alterations in brain dynamics associated with cognitive deficits in complex memory tasks. Crucially, these alterations occur prior to amyloid plaque onset, challenging the traditional amyloid-centric view of early AD pathology. Furthermore, we showed that two weeks of vGENUS increased brain dynamics between the first and the final day of the protocol. This increase in brain dynamics fluidity persisted after the end of stimulations, during performances of memory tasks where previously observed deficits were rescued.

Our findings suggest that alterations in awake-state brain dynamics could be an early hallmark of AD, occurring prior to amyloid plaque formation. The similarity between the hdEEG recordings in this study and those typically performed in humans suggests that brain dynamics fluidity could emerge as a promising diagnostic marker. This is supported by previous findings showing that microstates sequence complexity is reduced in AD patients and can help predict progression from Mild Cognitive Impairment (MCI) to AD (17). In addition, slowing down and increasing blocking frustration in resting state fMRI dynamic Functional Connectivity had already been reported in Alzheimer’s Disease (15) or cognitive challenge models of early mnesic impairments (42). Given the non-invasive nature and accessibility of EEG, assessing brain dynamics fluidity could enable earlier detection of AD, potentially during MCI and Subjective Cognitive Decline (SCD) stages, where patients report cognitive issues but show no deficits in standard neuropsychological tests. An easily implementable diagnostic tool for early-stage AD could significantly improve the timing and effectiveness of treatment interventions. The potential clinical translation of this approach warrants further investigation, including longitudinal studies in human populations to validate the predictive power of brain dynamics fluidity in AD progression.

Consistent with previous reports (21, 22), we found that 2 weeks of vGENUS rescued AD-related memory deficits in dKI mice. However, as recent studies have highlighted (27, 28), since 4-month-old dKI mice lack amyloid plaques, it is improbable that vGENUS targets amyloid pathology directly, or at least not exclusively or primarily. Instead, our data suggest that vGENUS restores healthy brain dynamics fluidity, an effect that persists even after stimulation ends. This finding points to a broader impact of vGENUS on brain dynamics beyond 40Hz entrainment, which has been debated in recent literature (26, 27). Notably, the restoration of global brain dynamics was not immediate but required a chronic 2-week protocol, indicating that long-term processes rather than immediate brain entrainment likely drive these changes.

One potential explanation for the observed effects could be metabolic reorganization of brain networks. Indeed, vGENUS has been shown to entrain vascular reactions in the brain, notably increasing blood vessel diameter (22, 25) and has already demonstrated beneficial effects in mouse models of stroke (43). Given that cerebral hypoperfusion has been linked to AD development (44) and cerebrovascular dysfunctions are associated with cognitive impairments (45), vGENUS-induced increases in brain blood flow could improve brain metabolism and, in turn, brain dynamics.

Another possibility is that vGENUS modulates specific neuromodulatory systems, particularly given that the reduction in brain dynamics fluidity observed in dKI mice is awake-state specific. vGENUS showed no effect on brain dynamics during sleep, suggesting that the mechanism may involve specific wake-related neuromodulators such as the noradrenergic system, which is both an early target in AD pathology (46) and known to affect large-scale brain dynamics (47). Future research will be needed to clarify the precise mechanisms by which vGENUS exerts its effects. However, our results, along with previous findings, converge to suggest that vGENUS beneficial effects on global brain dynamics are not specific to AD, as it has also shown benefits in epileptic patients (30). This suggests that vGENUS may represent a promising non-invasive treatment option for a variety of neurological disorders by targeting and modulating large-scale brain dynamics. Specifically, it could trigger endogeneous mechanisms for compensating early circuit dysfunction via a “reprogramming” of the working point of dynamic operation of brain networks (48, 49).

While our study provides novel insights into early AD pathophysiology and potential interventions, it is important to keep in mind that the use of a mouse model, while allowing for precise control and manipulation, may not fully recapitulate human AD pathology. Additionally, the long-term effects of vGENUS beyond the two-week period studied here remain to be explored.

In conclusion, our study reveals altered brain dynamics as an early marker of AD-related changes, detectable before significant amyloid plaque formation and correlating with subtle cognitive deficits. We demonstrate that vGENUS can restore these altered dynamics and ameliorate associated memory impairments. These findings not only provide new insights into early AD pathophysiology but also suggest novel approaches for early diagnosis and intervention in AD and potentially other neurological disorders characterized by disrupted brain dynamics.

## Materials and Methods

### Animals and surgery

*Subjects.* Double Knock-in App^NL-F^/MAPT (dKI) and Wild-Type (WT) mice were obtained as described in (32). For EEG-behavior experiments, 8 WT and 8 dKI mice were housed in individual cages post-surgery. For behavior only experiments, 35 WT and 37 dKI 4 months-old mice were housed in individual cages. All animals were under a 12h light/dark cycle with food and water present *ad libitum*. Both sexes were balanced in population to get closer to real population representation without studying sex effect. All experimental protocols agreed with the European Committee Council directive (2010/63/UE) and were approved by the French Ministry of Research (APAFIS#28839-2021010509459441).

*Surgery.* Surgeries were performed at 3 months old. Animals were anesthetized with isoflurane (IsoFlo, Zoetis) during the entire surgery (4% for induction then maintained at 1.5%). Local anesthesia was performed on incision site by Bupivacaïne and Lidocaïne (Lurocaine, Vetoquinol) injection prior to incision. Post-surgery analgesia was provided via sub cutaneous Metacam injection. EEG surface grid (H32 mouse EEG grid, Neuronexus, Ann Arbor, USA) were placed on the skull aligning the skull bregma with the grid landmark. The grid is fixed to the skull by applying Saline solution and letting it dry. One screw was inserted above the right cerebellum to serve as ground, and another one, placed rostral to the grid was used as fixation support for the implant. The implant was secured for long term use using dental glue (Super-bond, Sun Medical) and dental cement (Paladur).

### Mice perfusion and immunochemistry

Mice were deeply anesthetized with an intraperitoneal injection of ketamine (200 mg/kg) and xylazine (30 mg/kg) and then transcardially perfused with 0.1% heparin in 0.1 M phosphate-buffered saline (PBS), followed by 4% paraformaldehyde (PFA) in 0.1 M phosphate buffer (PB; pH 7.4, 4 °C). Brains were post-fixed in PFA for 24 hours, cryoprotected in 20% sucrose in PB for 48 hours, and subsequently stored at -80 °C. Coronal sections (40 µm) were obtained using a cryostat, covering the region from Bregma +2.58 to Bregma -5.88. Labeling was performed on one section every 160 µm, yielding approximately 62 sections per animal. Sections were processed at room temperature with agitation at 260 rpm as follows: three washes with PBS, followed by a 15-minute incubation in 70% formic acid, another three PBS washes, a 30-minute incubation in methanol with 0.3% H₂O₂, a 15-minute wash in ultrapure water, and two additional PBS washes. Sections were blocked for one hour in 5% normal horse serum (NHS) diluted in PBS containing 0.5% Triton X- 100, followed by an 18-hour incubation at 4°C with mouse anti-6E10 antibody (1:1000 in 2% NHS, BioLegend #803001). After three PBS washes, sections were incubated for two hours with a biotinylated horse anti-mouse antibody (1:500 in PBS containing 0.5% Triton X-100, Vector Laboratories, #BA-2001-.5), followed by three PBS washes. Sections were then incubated for 30 minutes in an avidin-biotin solution (Vector Laboratories), washed twice in PBS, and finally in PB- Tris. Detection was performed using a 15-minute incubation in 3,3-diaminobenzidine (Vector Laboratories). Sections were mounted onto gelatin-coated slides, air-dried for 24 hours, dehydrated through a graded series of alcohol baths (70%, 90%, 95%, 100%, 100%), cleared with Clearify (American MasterTech Scientific), and fixed on microscopic slides with Diamount (Diapath S.P.A). Slides were then dried for 48 hours in the dark. Whole-section images were acquired at 20x magnification using a Hamamatsu NanoZoomer S60 digital slide scanner (Hamamatsu Photonics K.K., Japan). Amyloid plaques were counted using the cell counter tool in ndpView2 software (Hamamatsu Photonics K.K., Japan). Plaques were identified based on their shape and the density of 6E10 labeling relative to the background. Specificity of labeling was confirmed by including a negative control in which the primary antibody (6E10) was omitted. As a positive control, sections from two 12-month-old dKI mice were processed and analyzed using the same protocol.

### Behavior

*Apparatus.* Behavioral tasks are conducted in a 55cm x 55cm open field with black wall and a white ground with a grid pattern.

*Habituation.* Two weeks after surgery, animals were habituated to the experimental apparatus. Transportation box was progressively presented over a week for the transport habituation. A recording cable was plugged several hours per day for 2 days to the head-mounted pre-amplifier so the mice could get used to its presence and weight. During the three days prior the behavioral task, an Open Field habituation consisting in ten minutes exploration of an empty Open Field and two object habituations consisting in two 10-minute sessions with a single object in the open field were performed.

*Novel Object Recognition Task.* At 4 months old and after vGENUS protocol, after the previously described habituation, mice perform a Novel Object Recognition (NOR) task. The mouse explores the open field where two similar objects are placed during a 10-minute sampling phase, and 24h after, realizing a 10-minute test phase consisting in an exploration of the open field where one of the previous objects is replaced by a new one. Objects exploration time is measured during the sampling and the test phase. To avoid place preference the object replaced is the one that was the less explored during the sampling phase.

*Object in Place Task.* At 4 months old and after vGENUS protocol, after the previously described NOR task, mice perform an Object in Place task. The mouse explores the open field where two different objects are placed during a 10-minute sampling phase, then wait a 5 minute inter trial interval in the home cage, before a second 10-minute test phase consisting in an exploration of the open field where one of the previous object is replaced by the copy of the other. Here neither the environment nor the object is new during the test phase but only the association between the specific object and its place in the environment. Objects exploration time is measured during the sampling and the test phase. To avoid place preference the object replaced is the one that was the less explored during the sampling phase.

*Memory performances.* To assess memory performances a Memory Index (MI) is computed from object exploration times during test as follow:

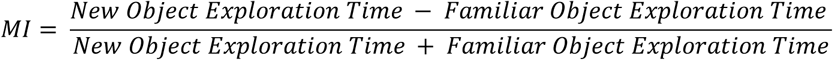

*Sleep recordings.* After the sampling phase of the NOR, mice are recorded in their home cage for a minimum of 6 hour during light phase to record first post learning sleep episodes.

### Visual Gamma Entrainment Stimulation (vGENUS) protocol

Visual GENUS protocol and setup was performed as described in (20). The day after the performance of the OiP task start a 15-day protocol. Each day, mice are placed in their home cages in the stimulation room with light on for a 1-hour stalling. Mice are then individually placed in stimulation cages, consisting in regular cages with 3 black painted walls to let transparent only the wall facing the stimulation light, without litter, food and water access. For one hour, the light of the room is turned off and only a 12V LED strip with a 4000/4500K light temperature and a 1200lm/m (Ledkia) controlled by an Arduino is either flickering at 40 Hz or displaying continuous light. After the stimulation, mice are back in the home cage for a 1-hour stalling in the stimulation room with the light on and the LED strip off before returning to the animal facility or doing behavioral task.

### Electrophysiological recordings and analysis

*Recording and preprocessing.* The electrophysiological activity was recorded with an Intan recording controller (RHD Recording Controller, Intan Technologies, USA). The signals were amplified 200x, recorded whole-band (0.1Hz-15 kHz) and digitized at 1kHz. Qualitative rejection of faulty channels was conducted, any animals presenting more than five faulty channels were excluded from the study. EEG signals were then lowpass filtered at 100Hz and artefact related activity were removed via Independent Component Analysis (ICA). The signals were then coarse-grained with a 40ms time windows to fit the frame rate of the behavior video recording.

*Spectral analysis.* Spectrograms were computed on non coarsegrained signals of 10 min recordings during vGENUS stimulation using Matlab Chronux toolbox *mtspecgram* function. Power spectrums were obtained by averaging the spectrogram over time. Relative 40Hz power was obtain by dividing the 40Hz power by the total power spectrum. 40Hz relative power was then averaged over cortical regions (Prefrontal Cortex:PFC; Motor Cortex: MOC; Somato-sensory Cortex: SSC; Parietal Cortex: PAR; Retrosplenial Cortex: RSC; Visual Cortex: VIS) based on Allen Institute mouse cortical parcelation.

*Microstates extraction with k-mean clustering.* Microstates were extracted independently for each animal and each trial to favorize individual characterization. To extract microstates sequences in an unsupervised way, k-mean clustering using native Matlab function with correlation distance were performed on the coarse-grained EEG time series for a wide ranges of cluster extraction (3 to 8).

*Microstates extraction with Global Field Power peaks.* To extract EEG microstates through a procedure closer to the human EEG literature (39), Global Field Power was computed as the variance of the coarse-grained EEG of all channels at each time points. Peak of GFP were determined using a minimum peak prominence of 0.5. EEG at GFP peaks was then clustered using k-mean clustering with native Matlab function with correlation distance and centroid were extracted. Microstates sequences were constructed by applying at each time point of the coarse-grained EEG the centroid showing the highest Pearson correlation.

*Microstates Fluidity.* To study switching across microstates, transition probability matrices were computed as the probability to switch from a microstate visited at time *t* to a different microstate at times *t*+1. Diagonal entries in this transition matrix correspond to the probability of remaining in the currently visited microstate without switching, which is proportional to the average dwell-time within the considered microstate. The average of diagonal transition matrix entities was thus a measure of microstates stability, and we defined its inverse as Microstates Fluidity. Such a quantity decreases if dwell-times increase and time to the next microstate switching is increased.

*Microstates Sequence Complexity.* Microstates sequence complexity is computed following (50), through a minimum description length approach. Following Kolmogorov and Chaitin (51), the shortest the symbolic sequence can be described in a suitable lossless-compressed format, the less complex it is. Each microstates sequence is considered as a sequence of symbolic labels corresponding to the microstate visited at each time. Let suppose that ***A***, ***B***, ***C****…* are the labels of different microstates. Their set forms the dictionary Δ. An original description of the sequence can be given by listing all the individual positions in the sequence where a given label is used, e.g. ***A*** *a1 a2 a3 …**B** b1 b2 b3 …**C** c1 c2 c3…* where *ai, bi, ci* …are positions where the labels ***A, B, C***… respectively appear in the sequence. The length of the original description is thus given by

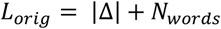

where |*Δ*| is the size of the dictionary and *Nwords* is the number of labels in the symbolic stream. If there are repetitions of consecutive symbols associated to permanence within a same microstate for a certain epoch of longer duration than a single time-step, a potentially compressed description of the same sequence can be obtained using a different encoding, ***A*** *l1 s1 l2 s2…**B** l1 s1 l2 s2…**C** l1 s1 l2 s2…* where the numbers *li* and *si* occurring alternatingly after a symbolic label, e.g. ***A,*** correspond, respectively to the length *li* of the *i*-th block of consecutive symbols ***A*** and to the shift of positions to find within the sequence the next (*i+1)-*th block of symbols ***A***. If the number of blocks of symbols is much smaller than the number of individual symbols in the sequence, then the length *L*_*comp*_ of this second description will be shorter than *L*_*orig*_. The complexity of the state transitions was quantified as the ratio of the compressed length to the original length Specifically, we also applied a base 100 exponential transformation so that the obtained values follow a normal distribution and statistical testing is simplified:

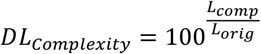

*Brain dynamics Fluidity.* What we call here Dynamics fluidity corresponds to a quantity which has been called inverse persistence in the literature about the analysis of dynamical systems and, specifically, climatic time-series (36). This quantity is evaluated based on empirical multivariate and continuous-valued time-series, here the 30-dimensional coarse-grained multi-channel EEG. The mathematical theory of this quantity is sophisticated, and we invite the reader to refer to the original publications for full detail (37, 52, 53). We can however provide some intuition. We consider the values *u*_*t,i*_ of the EEG activity of different channels *i* at time *t* as describing a trajectory ***ut*** in a high dimensional space. We can then consider a system configuration ***u0*** visited by the system at a certain time and ask how much time is needed for the system’s trajectory leaving a neighborhood of the current configuration, i.e. flowing away to far configurations that do not resemble ***u0***. The fluidity of the dynamics in proximity of the configuration ***u*** will be inversely reflecting the persistence time. The more persistent the configuration ***u***, the longer the previous and subsequent states of the system will resemble ***u***, and the lower will be the dynamics fluidity. To estimate fluidity from time-series observations, a connection between dynamical systems theory and extreme-value statistics can be exploited. To quantify the deviation of the trajectory ut from the reference point *u0*, we introduce a distance measure dist(*ut,u0*), using the Euclidean distance. We are interested in detecting clustering behavior, i.e., “sticky” behavior where the trajectory ut spends more time than usual around the reference point u0. To highlight the short distance values in these clustering situations, we use the log-distance and define the function *g*(*t;u0*)=−log(dist(*ut,u0*)). This function is large when the trajectory *ut* is in close proximity to *u0*. Consequently, the probability of persistence or returns around u0 corresponds to the probability of observing extreme values of g(t;u0).

Requiring that a point on the orbit falls within a ball of radius e^−*s*^ around *u0* is equivalent to asking that the corresponding value of the series *g*(*t*) exceeds the threshold s. Assuming independence of exceedances *g*(*t*), we obtain:

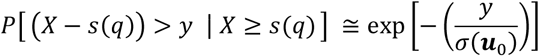

where *s(q)* is a high threshold associated with a quantile *q* of the series X≡g(t) The resulting distribution is an exponential member of the Generalized Pareto Distribution (GPD) family. The parameter σ, the scale parameter of the distribution, depends on the point *u0* in phase space and, for finite time series, on *t*. If we define *Mn* = max(*X0,X1,…,Xn−1),* we can then write:

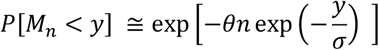

where *Mn* is the maximum of the series over *n* time-steps.The details of this computation are given in (54). A metric of persistence can then be obtained as the inverse of the extremal index θ, dimensionalized by the time step of the data used, whereas σ can be linked to the dimensionality of the data.

The parameters θ, μ and σ can be fitted practically on data using a suitable maximum likelihood estimator, by adopting a large threshold η. MATLAB code is provided in the Supporting Material for their evaluation, based on the Süveges estimator (55). In our study we set η to be equal to the 98% quantile of the distribution of *g(t; **u**0).* The quantity that we call here dynamics fluidity corresponds to the fitted parameter θ. The larger will be θ, the less persistent (and thus the more fluid) will be the dynamics. The case of θ = 0 corresponds notably to ***u****0* being a fixed point, so that the probability of remaining in its proximity after having reached it is 1. The case of θ = 1 corresponds to the probability of the return time being a Poisson distribution and thus to the absence of trajectory points clustering. Furthermore, the parameter σ provides information on the local dimension of the attractor manifold surrounding ***u****0* (see again (36)) but we do not exploit this information in this study.

The estimated parameters, including dynamics fluidity θ depend on the local point *u*_0_ chosen as reference. Thus, fluidity analysis of a multivariate time-series of EEG will yield a time-series of values of fluidity, each estimated using as reference a different instantaneous observation of EEG multi-channel topography.

### Statistical analysis

All statistics were performed using either built-in Matlab (r2024a) functions, Toolboxes, custom script or Jamovi 2.5. For all statistical test the significance threshold α was fixed at 0.05.

*Behavior.* Memory Index was analyzed using either unpaired t-test between both genotypes for each task before and after vGENUS for implanted animals displayed in Fig.1 & 5, 2-way ANOVA (factor: Genotype, Task) were used on the larger non implanted cohort before vGENUS and repeated 2-way ANOVA (non repeated factor: Genotype; repeated factor: vGENUS) were used on the larger non implanted cohort after vGENUS. Post hoc tests were used when necessary, using a Bonferroni correction for multiple comparison. Before and after GENUS, memory index was compared for each Genotypes against chance levels using one sided t-test testing H1 > 0 or memory index “above” chance levels. Exploration Time was analyzed using a 3-way ANOVA (factor: Genotype, Task and Phase (Sampling/Test) before vGENUS and 3-way repeated ANOVA (non-repeated factor: Genotype and Phase (Sampling/Test); repeated factor: GENUS (before/after)) after vGENUS. Post hoc tests were used when necessary, using a Bonferroni correction for multiple comparison.

*EEG Microstates.* Microstates Fluidity and Complexity were analyzed using 4-way ANOVA (factor: Genotype, Phase, Clusters, Task; SI Appendix Fig. S2). Post hoc tests were used when necessary, using a Bonferroni correction for multiple comparison. As the factor Phase (Sampling/Test) & Clusters showed no significant interaction with other factors, to reduce dimensions analyses were further conducted on concatenated EEG between Sampling and Phase, and Microstates fluidity and complexity averaged over the number of microstates extracted with a 2-way ANOVA (factor: Genotype, Task; Fig. 2). Beside similar 2-way ANOVA (Fig. 5), post vGENUS results were also analyzed with a repeated 4-way ANOVA (factor: Genotype, Task, Clusters, vGENUS, SI Appendix Fig. 5).

*Dynamics Fluidity.* EEG fluidity was computed on EEG time-series concatenated between Sampling and Test. Average distributions were then compared two by two using bootstrapped Kolmogorov-Smirnoff (KS) statistics to probe whether the two distributions had significantly shifted mean and range respectively to a null hypothesis of no shift.

Using a Montecarlo procedure (pre-implemented in Matlab via the randsample function with a controlled random number stream), we redrawn 5000 random samples according to each of the distributions of fluidity to compare (modeled as histograms with a resolution of 200 uniform-width bins, with different histograms for different genotypes and conditions before of after vGENUS). We performed then random subsampling of these large Montecarlo samples, generating reduced bootstrap with replacements replicas each including 500 resampled observations, under two alternative hypotheses. In a H1 hypothesis of association between fluidity and genotype and condition, subsampling was performed separately over each of the Montecarlo sample specific to the different cases. In a H0 hypothesis of lack of association, we merged the Montecarlo samples for the two genotypes and generated two random subsamples from this common merged sample. We then computed KS statistics between the two subsamples, quantifying over bootstrap replicas the distribution of KS statistics between fluidity distributions for different genotypes, under both the H1 and H0 hypotheses. In the Results, we communicate the mean and standard deviation of KS statistics across iterations in the association hypothesis. Statistical significance of the difference between genotypes was assessed by comparing the H1-distribution of KS divergence values with the chance-level distribution under the H0 hypotheses, estimating a *p*-value based on the fraction of overlap between the two distributions, and correctd for multiple comparison with a Bonferroni correction. Note that this procedure is more conservative than the classical testing based on KS statistics non-parametric comparison.

*Continuous dynamics-Microstates fluidity correlation.* For each animal OiP and NOR preGENUS, mean dynamics fluidity was computed as the average of the EEG dynamics fluidity previously computed, mean microstates fluidity and complexity were computed by averaging microstates fluidity or complexity obtained with 3 to 8 clusters to obtain a single value. Pearson correlations were then computed between mean dynamics fluidity and mean microstates fluidity or mean dynamics fluidity and mean microstates complexity taking all animals during OiP and NOR.

## Acknowledgments

This work was supported by CNRS, Université de Strasbourg and grants from Agence Nationale de la Recherche: ANR HippoComp (ANR-21-CE37-0011) to RG and DB, and Fondation pour la Recherche Médicale (FRM): FRM- ALZ201912009643 to CM. The authors wish to thank Pr. Jesse Jackson and Dr. Yaroslav Sych for critical reading of the manuscript, Viktor Jirsa for inspiring discussions, the PIV (Plateforme d’Imagerie in Vitro, INCI, Strasbourg), Coline Portet and Flora Thellier, PhD students from the lab, for offering their technical help for the experiments.

## Author Contributions

M.A. and R.G. performed the experiments and analyzed the data. C.M. provided the animals and gave inputs for behavioral experiments. K.H. performed immunochemistry experiments. A.I. managed the animal breeding and genotyping. D.F. provided code for dynamic fluidity analyses. R.G. and D.B. conceived the experimental and analytical design; M.A., R.G. and D.B. wrote the article.

