## Supporting Information for "40 Hz light stimulation restores early brain dynamics alterations and associative memory in Alzheimer’s disease model mice"

**This PDF file includes:**

Figures S1 to S6

**Supporting Information Text**

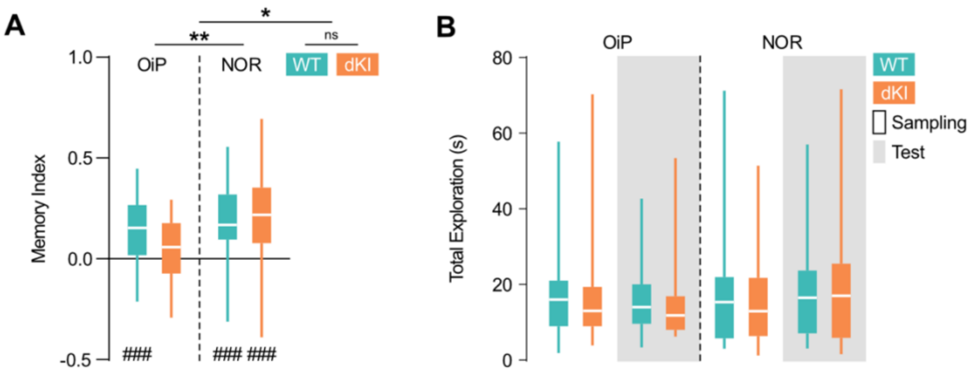


**Fig. S1. dKI mice memory deficits in complex task are unrelated to exploration time.** Box ranges from 25 to 75 percentile and whiskers for minimum to maximum values, median is represented by white line. **(A)** Memory Index for WT (blue, n = 35) and dKI (orange, n = 37) mices performing Object in Place (OiP) and Novel Object Recognition (NOR) Tasks. Two way ANOVA (factor: Task, Genotype) reveal a significant interaction between Genotype and Task ($F(1,140) = 5.17$, $p = 0.024$) and only dKI mice performing OiP showed performances not higher than chance level (one sided t-test against chance, # : $p < 0.05$, ### : $p < 0.001$). **(B)** Total exploration time of the two objects for WT and dKI mice during Sampling (no background) and Test (grey background) of OiP and NOR task. Three way ANOVA (factor: Genotype, Task, Phase) shows no significant effects, thus indicating no difference in exploration between genotypes in both tasks.

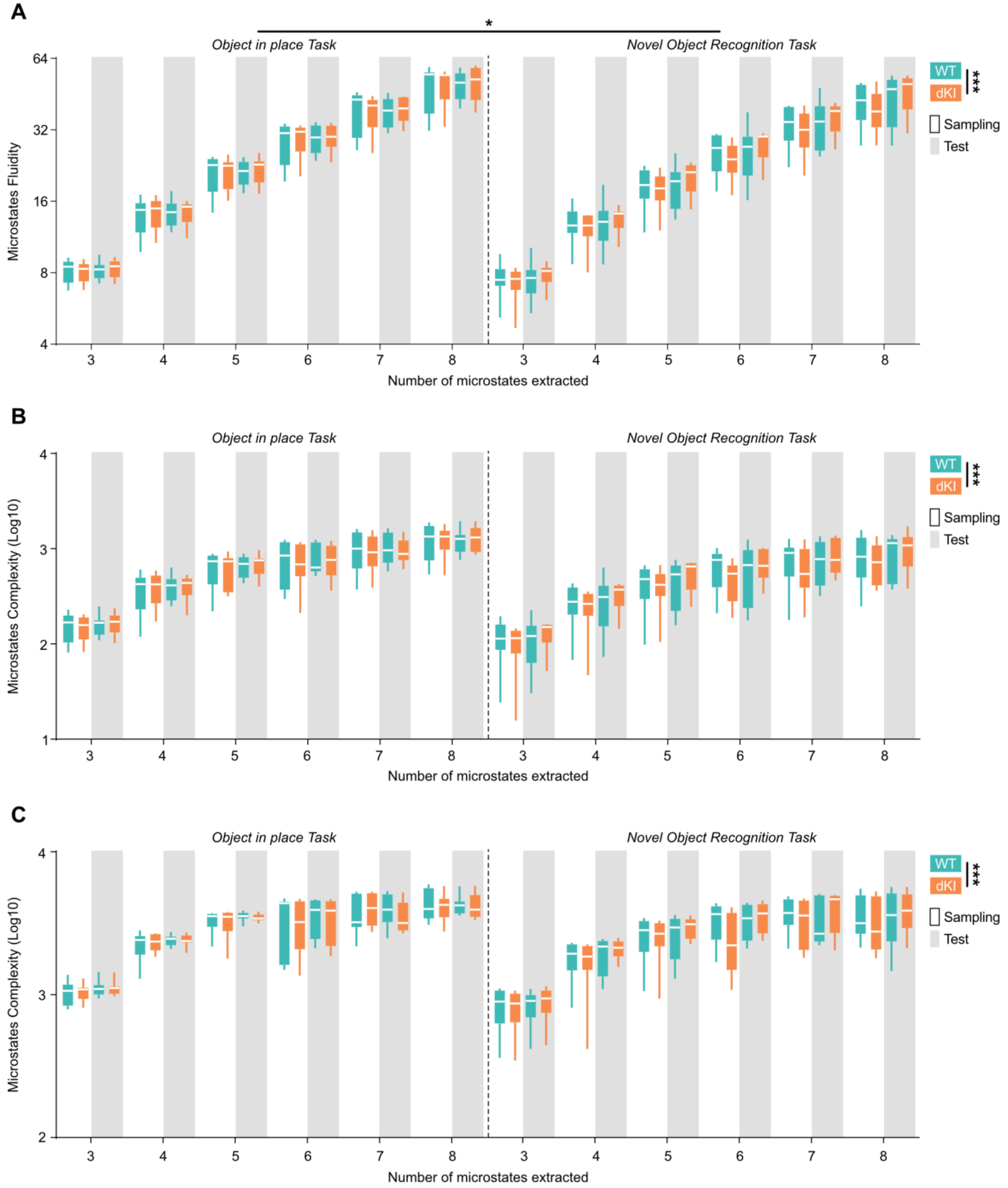
A
*
Object in place Task
Novel Object Recognition Task
Microstates Fluidity
64
32
16
8
4
WT
dKI
***
Sampling
Test
3 4 5 6 7 8 3 4 5 6 7 8
Number of microstates extracted
B
Object in place Task
Novel Object Recognition Task
Microstates Complexity (Log10)
4
3
2
1
WT
dKI
***
Sampling
Test
3 4 5 6 7 8 3 4 5 6 7 8
Number of microstates extracted
C
Object in place Task
Novel Object Recognition Task
Microstates Complexity (Log10)
4
3
2
WT
dKI
***
Sampling
Test
3 4 5 6 7 8 3 4 5 6 7 8
Number of microstates extracted

**Fig. S2. Microstates dynamics and complexity alteration are independent from the number of microstates extracted and not only explained by microstates repetition.** Box ranges from 25 to 75 percentile and whiskers for minimum to maximum values, median is represented by white line. **(A)** Microstate fluidity during sampling (no background) and test (grey background) phase of OiP and NOR tasks for WT (blue) and dKI (orange) mice (n=8 per group) across 3 to 8 microstates extracted. Four-way ANOVA (factor: Genotype, Task, Phase, Clusters; *: $p<0.05$ **: $p<0.01$ ***: $p<0.001$) showed significant Genotype effect ($F(1,336) = 41.884$; $p < 0.001$) indicating a lower fluidity in dKI mice. This effect showed no interactions with Task or Phase. **(B)** Microstate sequence complexity during sampling (no background) and test (grey background) phase of OiP and NOR tasks for WT (blue) and dKI (orange) mice (n=8 per group) across 3 to 8 microstates extracted. Four-way ANOVA (factor: Genotype, Task, Phase, Clusters; *: $p<0.05$ **: $p<0.01$ ***: $p<0.001$) showed significant Genotype effect ($F(1,336) = 40.185$; $p < 0.001$) indicating a lower complexity in dKI mice. This effect showed no interactions with Task or Phase. **(C)** Repetition free microstate sequence complexity during sampling (no background) and test (grey background) phase of OiP and NOR tasks for WT (blue) and dKI (orange) mice (n=8 per group) across 3 to 8 microstates extracted. Four-way ANOVA (factor: Genotype, Task, Phase, Clusters; *: $p<0.05$ **: $p<0.01$ ***: $p<0.001$) showed significant Genotype effect ($F(1,336) = 22.701$; $p < 0.001$) indicating a lower complexity in dKI mice. This effect showed no interactions with Task or Phase.

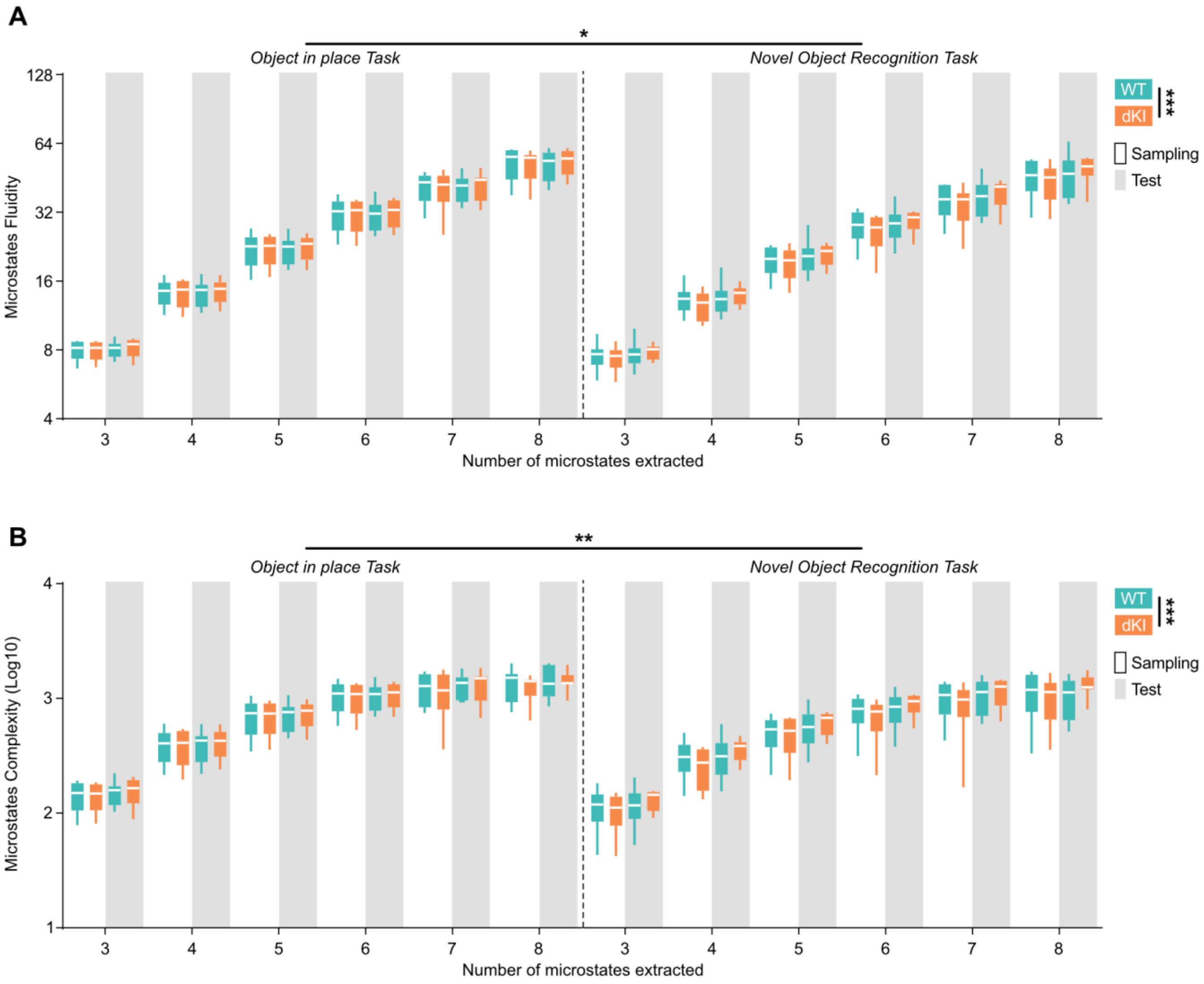


**Fig. S3. Microstates sequence dynamics and complexity are altered in dKI mice and rescued by vGENUS when using Global Field Power microstates extraction method.** Box ranges from 25 to 75 percentile and whiskers for minimum to maximum values, median is represented by white line. **(A)** Microstate fluidity during sampling (no background) and test (grey background) phase of OiP and NOR tasks for WT (blue) and dKI (orange) mice (n=8 per group) across 3 to 8 microstates extracted. Four-way ANOVA (factor: Genotype, Task, Phase, Clusters; *: $p<0.05$ **: $p<0.01$ ***: $p<0.001$) showed significant Genotype effect ($F(1,336) = 33.948$; $p < 0.001$) indicating a lower fluidity in dKI mice. This effect showed no interactions with Task or Phase. **(B)** Microstate sequence complexity during sampling (no background) and test (grey background) phase of OiP and NOR tasks for WT (blue) and dKI (orange) mice (n=8 per group) across 3 to 8 microstates extracted. Four-way ANOVA (factor: Genotype, Task, Phase, Clusters; *: $p<0.05$ **: $p<0.01$ ***: $p<0.001$)

showed significant Genotype effect ($F(1,336) = 39.279$; $p < 0.001$) indicating a lower complexity in dKI mice. This effect showed no interactions with Task or Phase.

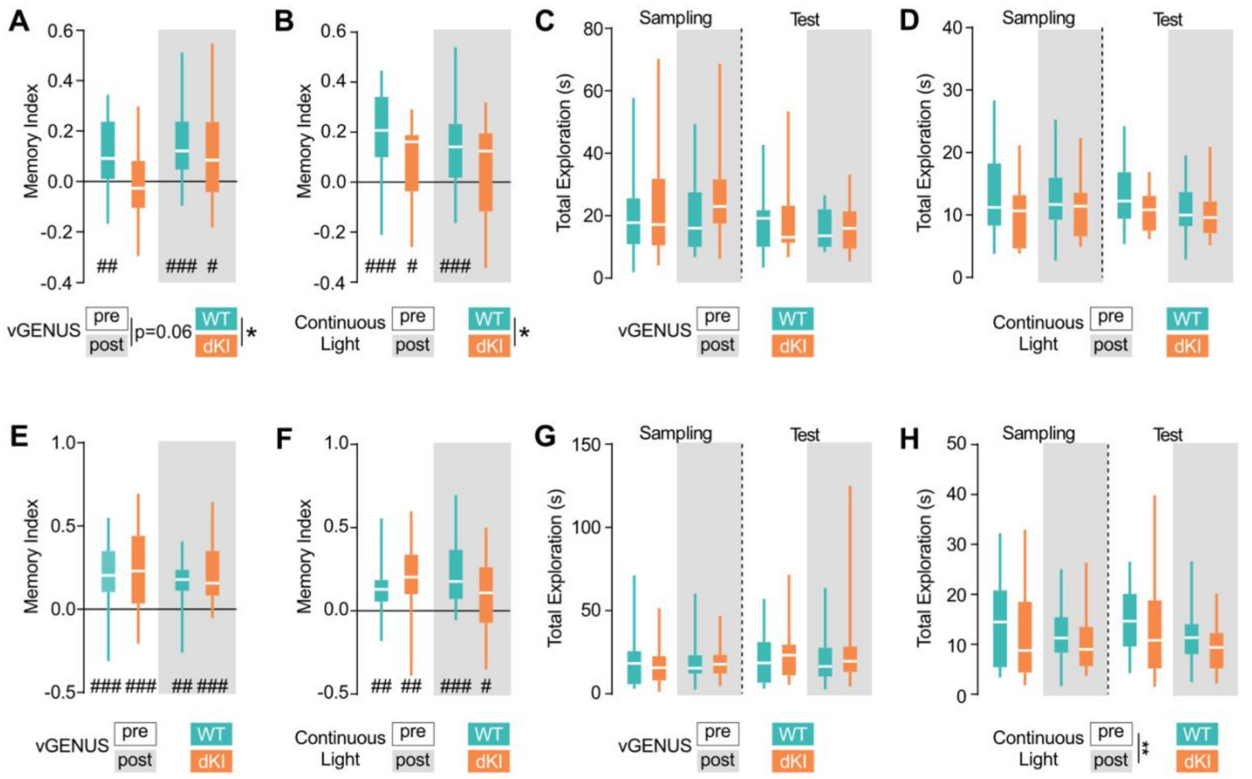


**Fig. S4. vGENUS restoration of memory performances is dependant on 40Hz frequency.** Box ranges from 25 to 75 percentile and whiskers for minimum to maximum values, median is represented by white line. **(A)** Memory index for the OiP task in WT (n=17, blue) and dKI (n=19, orange) non EEG-recorded mice pre- (no background) and post- (grey background) vGENUS. Two-way ANOVA on repeated measures showed a genotype effect ($F(1,34) = 4.46$, $p = 0.042$) but no stimulation effect ($F(1,34) = 3.736$, $p = 0.062$). In dKI mice, memory performance did not exceed chance levels pre-vGENUS but improved significantly post-vGENUS (one-sided t-test against chance, #: $p < 0.05$, ##: $p < 0.01$, ###: $p < 0.001$). **(B)** Memory index for the OiP task in WT (n=18, blue) and dKI (n=18, orange) non EEG-recorded mice pre- (no background) and post- (grey background) 2 weeks of daily continuous light stimulation. Two-way ANOVA on repeated measures showed a genotype effect ($F(1,34) = 5.38$, $p = 0.026$) but no stimulation effect ($F(1,34) = 0.4463$, $p = 0.509$). dKI mice showed a memory index not higher from chance chance post-Continuous light (one-sided t-test against chance, #: $p < 0.05$, ##: $p < 0.01$, ###: $p < 0.001$). **(C)** Total exploration time during OiP task in WT (n=17, blue) and dKI (n=19, orange) non EEG-recorded mice pre- (no background) and post- (grey background) vGENUS. Three-way ANOVA on repeated measures showed no genotype ($F(1,68) = 1.208$, $p = 0.276$), vGENUS ($F(1,68) = 0.411$, $p = 0.524$), or phase effects ($F(1,68) = 2.953$, $p = 0.09$). **(D)** Total exploration time during OiP task in WT (n=18, blue) and dKI (n=18, orange) non EEG-recorded mice pre- (no background) and post- (grey background)

2 weeks of daily continuous light stimulation. Three-way ANOVA on repeated measures showed no genotype ($F(1,68) = 2.901$, $p = 0.093$), stimulation ($F(1,68) = 1.324$, $p = 0.254$), or phase effect ($F(1,68) = 0.714$, $p = 0.401$). **(E)** Memory index for the NOR task in WT (n=17, blue) and dKI (n=19, orange) non EEG-recorded mice pre- (no background) and post- (grey background) vGENUS. Two-way ANOVA on repeated measures showed no genotype ($F(1,34) = 0.698$, $p = 0.409$) nor vGENUS effect ($F(1,34) = 0.117$, $p = 0.734$). Both WT and dKI mice memory performances were higher than chance levels pre and post-vGENUS (one-sided t-test against chance, #: $p < 0.05$, ##: $p < 0.01$, ###: $p < 0.001$). **(F)** Memory index for the NOR task in WT (n=18, blue) and dKI (n=18, orange) non EEG-recorded mice pre- (no background) and post- (grey background) 2 weeks of daily continuous light stimulation. Two-way ANOVA on repeated measures showed no genotype ($F(1,34) = 0.449$, $p = 0.507$) nor stimulation effect ($F(1,34) = 0.218$, $p = 0.643$). Both WT and dKI mice memory performances were higher than chance levels pre and post-Continuous light (one-sided t-test against chance, #: $p < 0.05$, ##: $p < 0.01$, ###: $p < 0.001$). **(G)** Total exploration time during NOR task in WT (n=17, blue) and dKI (n=19, orange) non EEG-recorded mice pre- (no background) and post- (grey background) vGENUS. Three-way ANOVA on repeated measures showed no genotype ($F(1,68) = 0.648$, $p = 0.424$), vGENUS ($F(1,68) = 0.544$, $p = 0.463$), or phase effects ($F(1,68) = 1.403$, $p = 0.240$). **(H)** Total exploration time during NOR task in WT (n=18, blue) and dKI (n=18, orange) non EEG-recorded mice pre- (no background) and post- (grey background) 2 weeks of daily continuous light stimulation. Three-way ANOVA on repeated measures showed no genotype ($F(1,68) = 1.367$, $p = 0.246$) or phase effect ($F(1,68) = 9.78x10-4$, $p = 0.975$) but a stimulation effect ($F(1,68) = 11.08$, $p = 0.001$) indicating a reduced exploration after 2 weeks of continuous light exposure in both genotypes.

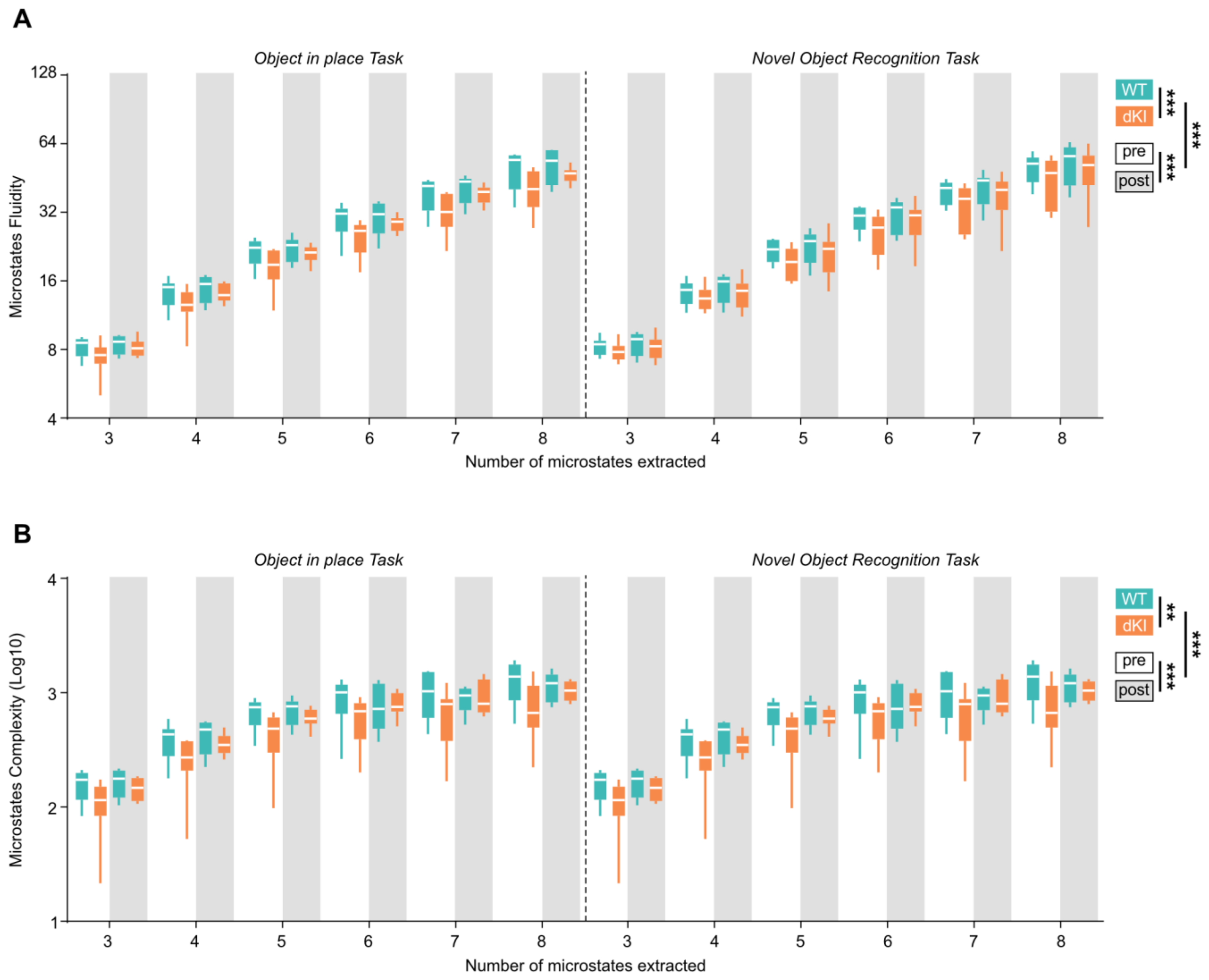


**Fig. S5. Microstates dynamics and complexity restoration by vGENUS are independent from the number of microstates extracted.** Box ranges from 25 to 75 percentile and whiskers for minimum to maximum values, median is represented by white line. **(A)** Microstate fluidity during OiP and NOR tasks for WT (blue) and dKI (orange) mice pre (no background) and post (grey background) vGENUS (n=8 per group) across 3 to 8 microstates extracted. Four-way repeated ANOVA (non –repeated factors: Genotype, Task, Clusters; repeated factor: vGENUS; *: $p<0.05$ **: $p<0.01$ ***: $p<0.001$) showed significant interaction between Genotype and vGENUS ($F(1,168) = 12.265$; $p < 0.001$). Post hoc test showed that vGENUS increased fluidity in WT ($t(168) = -3.09$; $p = 0.014$) and in dKI mice ($t(168) = -8.05$; $p <0.001$) bringing a fluidity lower than WT ($t(168) = 5.23$; $p < 0.001$) to a fluidity almost similar to WT level ($t(168) = 2.69$, $p = 0.048$). **(B)** Microstate sequence complexity during OiP and NOR tasks for WT (blue) and dKI (orange) mice pre (no background) and post (grey background) vGENUS (n=8 per group) across 3 to 8 microstates extracted. Four-way repeated ANOVA (non –repeated factors: Genotype, Task, Clusters; repeated factor: vGENUS; *: $p<0.05$ **: $p<0.01$ ***: $p<0.001$) showed significant interaction between Genotype and vGENUS ($F(1,168) = 15.865$; $p < 0.001$). Post hoc test showed that vGENUS increased fluidity in WT ($t(168) = -5.144$; $p < 0.001$) and in dKI mice ($t(168) = -4.869$; $p <0.001$) bringing a complexity lower than WT ($t(168) = 5.443$; $p < 0.001$) to a complexity non different to WT level ($t(168) = 2.153$, $p = 0.196$).

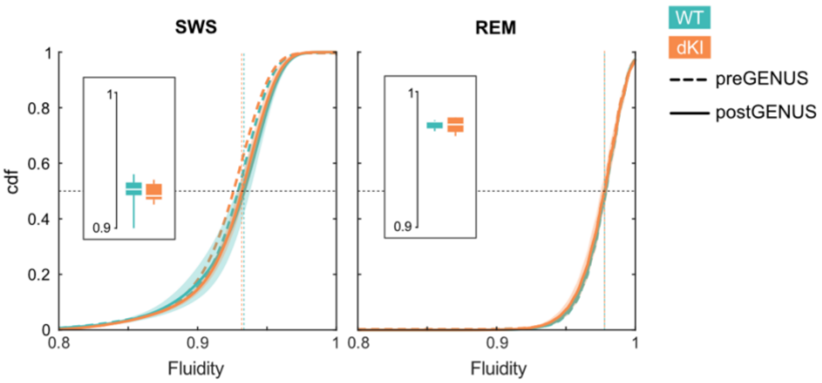


**Fig. S6. vGENUS induces no change in sleep brain fluidity.** Brain dynamics fluidity cumulative density distribution for WT and dKI mice show no differences in both REM and SWS after vGENUS (plain curve). No significant differences were observed for each genotype with dynamics fluidity before vGENUS (dotted curve). Data are presented as mean ± s.e.m. Dotted lines show distribution medians. Box displays individuals mean dynamics fluidity distributions.
